# Leaf anatomy is not correlated to CAM function in a C_3_+CAM hybrid species, *Yucca gloriosa*

**DOI:** 10.1101/726737

**Authors:** Karolina Heyduk, Jeremy N. Ray, Jim Leebens-Mack

**Affiliations:** Department of Ecology and Evolutionary Biology, Yale University, New Haven, CT USA; Department of Plant Biology, University of Georgia, Athens, GA USA

**Keywords:** *Yucca gloriosa*, *Yucca aloifolia*, *Yucca filamentosa*, Asparagaceae, Agavoideae, CAM photosynthesis, leaf anatomy, hybrid

## Abstract

**Background and Aims:** CAM photosynthesis is often considered to be a complex trait, requiring orchestration of leaf anatomy and physiology for optimal performance. But the observation of trait correlations is based largely on comparisons between C_3_ and strong CAM species, resulting in a lack of understanding as to how such traits evolve and the level of intraspecific variability for CAM and associated traits.

**Methods:** To understand intraspecific variation for traits underlying CAM and how these traits might assemble over evolutionary time, we conducted detailed time course physiological screens and measured aspects of leaf anatomy in 24 genotypes of a C_3_+CAM hybrid species, *Yucca gloriosa* (Asparagaceae). Comparisons were made to *Y. gloriosas* progenitor species, *Y. filamentosa* (C_3_) and *Y. aloifolia* (CAM).

**Key results:** Based on gas exchange and measurement of leaf acids, *Y. gloriosa* appears to use both C_3_ and CAM, and varies across genotypes in the degree to which CAM can be upregulated under drought stress. While correlations between leaf anatomy and physiology exist when testing across all three *Yucca* species, such correlations break down at the species level in *Y. gloriosa*.

**Conclusions:** The variation in CAM upregulation in *Y. gloriosa* is a result of its relatively recent hybrid origin. The lack of trait correlations between anatomy and physiology within *Y. gloriosa* indicate that the evolution of CAM, at least initially, can proceed through a wide combination of traits, and more favorable combinations are eventually selected for in strong CAM plants.

## Introduction

A fundamental aim of comparative biology is to elucidate how, when, and why traits evolve, and the biological consequences of trait evolution. Some traits have simple genetic architecture: changes may be induced by mutations to single genes or regulatory elements, as in the case of hair color in mice (Hoekstra *et al*. 2006), flower color and pollinator shifts in *Erythranthe guttata* (Bradshaw and Schemske 2003; Yuan *et al*. 2013), and herbicide resistance in barley (Lee *et al*. 2011). Other traits are more complex, in that they are actually a sum of phenotypic states orchestrated across an organism. For example, the evolution of C_4_ photosynthesis requires both an altered biochemical pathways as well as changes to leaf vein density and anatomy (Hatch 1987; Christin *et al*. 2013; Sage *et al*. 2014), and burrowing behavior in field mice relies on changes to separate genetic modules (Weber *et al*. 2013). Complex traits are unlikely to evolve via a single mutation (Lenski *et al*. 2003), and as a result one might expect various intermediate phenotypes to exist through the evolutionary progression from ancestral to derived character states. Species exhibiting intermediate phenotypes could be instrumental to ordering the sequence of events that led to the evolution of a complex trait. Intermediate phenotypes also lend insight into the genetic landscape of a complex trait: for example, genetic linkage can restrict which trait combinations are possible and can affect how quickly natural selection can act upon them.

Crassulacean acid metabolism (CAM) is an example of a complex plant trait - involving biochemistry, anatomy, and physiology - whose evolutionary trajectory remains obscured, despite multiple independent origins (Edwards 2019). CAM is a carbon concentrating mechanism that works tandemly with the C_3_ Calvin Benson cycle to increase the water use efficiency of plants. The C_3_ pathway uses Rubisco, an enzyme which has both carboxylating and oxygenating functions. Under high temperatures or conditions that promote stomatal closure, rates of Rubisco oxygenation increase and force plants to undergo photorespiration, an energetically costly process that fixes no net CO_2_. CAM concentrates CO_2_ in an effort to reduce oxygenation via Rubisco and photorespiratory stress. CAM species open their stomata at night, when cooler temperatures and higher relative humidity reduce evapotranspiration. Incoming CO_2_ is initially converted to malate and stored in the vacuole. During the day, the stomata largely remain closed, and the stored CO_2_ is decarboxylated from malate, surrounding Rubisco and the C_3_ machinery with elevated levels of CO_2_. The carbon concentrating mechanism of CAM reduces levels of photorespiration while simultaneously increasing overall water use efficiency. As a result, CAM plants are often found in hot, arid, or seasonally dry habitats - often, but not always, where water is limiting.

Since all CAM plants retain and use the entire C_3_ machinery, many species fix carbon through a mixture of both pathways (Winter 2019). Strong CAM plants use CAM for the vast majority of their carbon uptake, while C_3_+CAM species use a mix of both pathways to fix CO_2_. CAM cycling plants fix respired CO_2_ nocturnally through the CAM pathway but otherwise have C_3_ physiology. Moreover, plants can vary not only in their ability to use CAM, but also the degree to which CAM can be modulated under abiotic stress. Both strong CAM and C_3_+CAM can alter the relative contribution of CAM to CO_2_ fixation as a response to abiotic stressors. C_3_+CAM species can up-regulate the CAM pathway (“facultative CAM”) or downregulate the contribution of the C_3_ pathway, whereas strong CAM species may increase the degree of C_3_ carbon fixation when exceptionally well-watered (Hartsock and Nobel 1976). It is unclear how intermediate phenotypes such as C_3_+CAM and CAM cycling fit into the evolutionary trajectory of CAM, but the prevalence of intermediate CAM species (Winter 2019) suggests that such a dynamic phenotype can be advantageous under certain situations (i.e., seasonal drought) (Winter *et al*. 2008; Herrera 2009).

CAM photosynthesis has evolved at least 60 times independently (Edwards and Ogburn 2012), though this is likely an inaccurate count due to the difficulties associated with surveying intermediate CAM, particularly facultative forms involving induction of CAM only under specific conditions. Additionally, attempts to delineate when CAM evolved within extant CAM lineages are made difficult by a lack of phylogenetic resolution, particularly in very diverse lineages. However, the repeated origin of CAM suggests that the transition to CAM from a C_3_ ancestor may be evolutionarily straightforward (Heyduk, Moreno-Villena, *et al*. 2019). Indeed, the entire CAM biochemical pathway is present in all C_3_ plants as a part of the tricarboxylic acid cycle, and thus all genes required for CAM are already present in all C_3_ angiosperms. Most, if not all, genes involved in coding the CAM biochemical reactions are found in multiple copies in angiosperm genomes, allowing a single member of a gene family to be recruited for CAM (Silvera *et al*. 2014; Ming *et al*. 2015; Yang *et al*. 2017). There is mixed evidence that amino acid substitutions are required for evolutionary shifts to CAM function (Yang *et al*. 2017; Goolsby *et al*. 2018), and many recent genomic studies have suggested that changes to regulation of gene transcription and translation are critical for CAM evolution (Heyduk *et al*. 2018; Yin *et al*. 2018; Wai *et al*. 2019).

In addition to the importance of expressing the right genes at the right time, specific anatomical traits have long thought to be required for maximum CAM function (Nelson *et al*. 2005; Nelson and Sage 2008). To be able to store large amounts of CO_2_ as malate, CAM plants require larger vacuoles; indeed CAM plants typically have larger mesophyll cells than their C_3_ counterparts (Heyduk, Burrell, *et al*. 2016; Males 2018). Intercellular airspace (IAS) is often reduced in CAM species (Nelson and Sage 2008; Zambrano *et al*. 2014). One theory is that tight packing of cells reduces the amount of CO_2_ leakage that can occur during the day, when malate is decarboxylated and results in high concentrations of CO_2_ in the cells. Alternatively, reduced IAS may just be a result of larger cells packed into a leaf whose size may be limited by other factors, including the need to maintain hydraulic connectivity. Finally, CAM plants are often described as having thick, succulent leaves. The importance and timing of these anatomical changes remains unclear: in some systems, species that are C_3_+CAM look anatomically like their C_3_ relatives (Silvera *et al*. 2005; Males 2018), whereas other lineages evolve succulent leaf anatomy prior to CAM (Heyduk, McKain, *et al*. 2016) or coincident with the origin of CAM (Zambrano *et al*. 2014). As a result, our understanding of the importance of leaf anatomy on CAM function remains unclear.

One of the greatest challenges in understanding the concerted evolution between CAM biochemistry and anatomy is a lack of systems in which genetic segregation produces variation within and among these traits. While comparisons between C_3_ and strong CAM species have helped us define a suite of traits that seem to segregate with photosynthetic pathway, these comparisons conflate trait differences with evolutionary distance and yield little insight into how suites of CAM traits have been assembled repeatedly in plant evolutionary history. Are traits assembled sequentially, such that a certain phenotype must arise (e.g., large cells) before a secondary phenotype can evolve (e.g., accumulation of malate)? Or are there a number of trait combinations that can arise in any order and span phenotypic space, but selection repeatedly favors certain combinations to maximize the efficiency of CAM? These questions can be addressed by studying intraspecific variation of anatomy and photosynthetic physiology in an intermediate C_3_+CAM species.

To understand how anatomy and physiology are correlated - or not - we measured anatomical and photosynthetic traits in a C_3_+CAM species, *Yucca gloriosa*, a naturally occurring homoploid hybrid species resulting from a wild cross between *Y. aloifolia* (CAM) and *Y. filamentosa* (C_3_) (Rentsch and Leebens-Mack 2012; Heyduk, Burrell, *et al*. 2016). All three *Yucca* species overlap in the southeastern United States, with *Y. filamentosa* having the broadest range that extends into the northeast and midwest, *Y. aloifolia* being more restricted to the southeast, and *Y. gloriosa* inhabiting the narrowest natural range, occurring only in the coastal regions of the Atlantic seaboard between Florida and Virginia. Previous work has demonstrated contrasting photosynthetic pathways in the parental species and intermediate physiology and anatomy in the hybrid (Heyduk, Burrell, *et al*. 2016; Heyduk, Ray, *et al*. 2019). Genetic screens based on microsatellite data have suggested that while *Y. gloriosa* still retains a mixture of both parental genomes, genotypes are not identical and thus not likely to be F1 hybrids (Rentsch and Leebens-Mack 2012; Heyduk, Burrell, *et al*. 2016). With genetic variation in mind, here we expand our previous sampling to measure gas exchange, leaf acid accumulation, and anatomical characteristics in 24 genotypes of *Y. gloriosa*. We tested whether 1) *Y. gloriosa* genotypes vary in physiological and anatomical traits related to CAM and in their response to drought stress, and 2) if anatomical traits are closely correlated with the photosynthetic phenotype of a genotype (Fig. 1). We show that genotypes vary in the degree of CAM used, as well as the level of upregulation of CAM under drought stress; we further find that there is little correlation between anatomical traits and photosynthetic phenotypes. The lack of trait correlations within a species suggest the processes that shape trait distributions between species are uncoupled from those occurring within a species.

**Figure 1.**
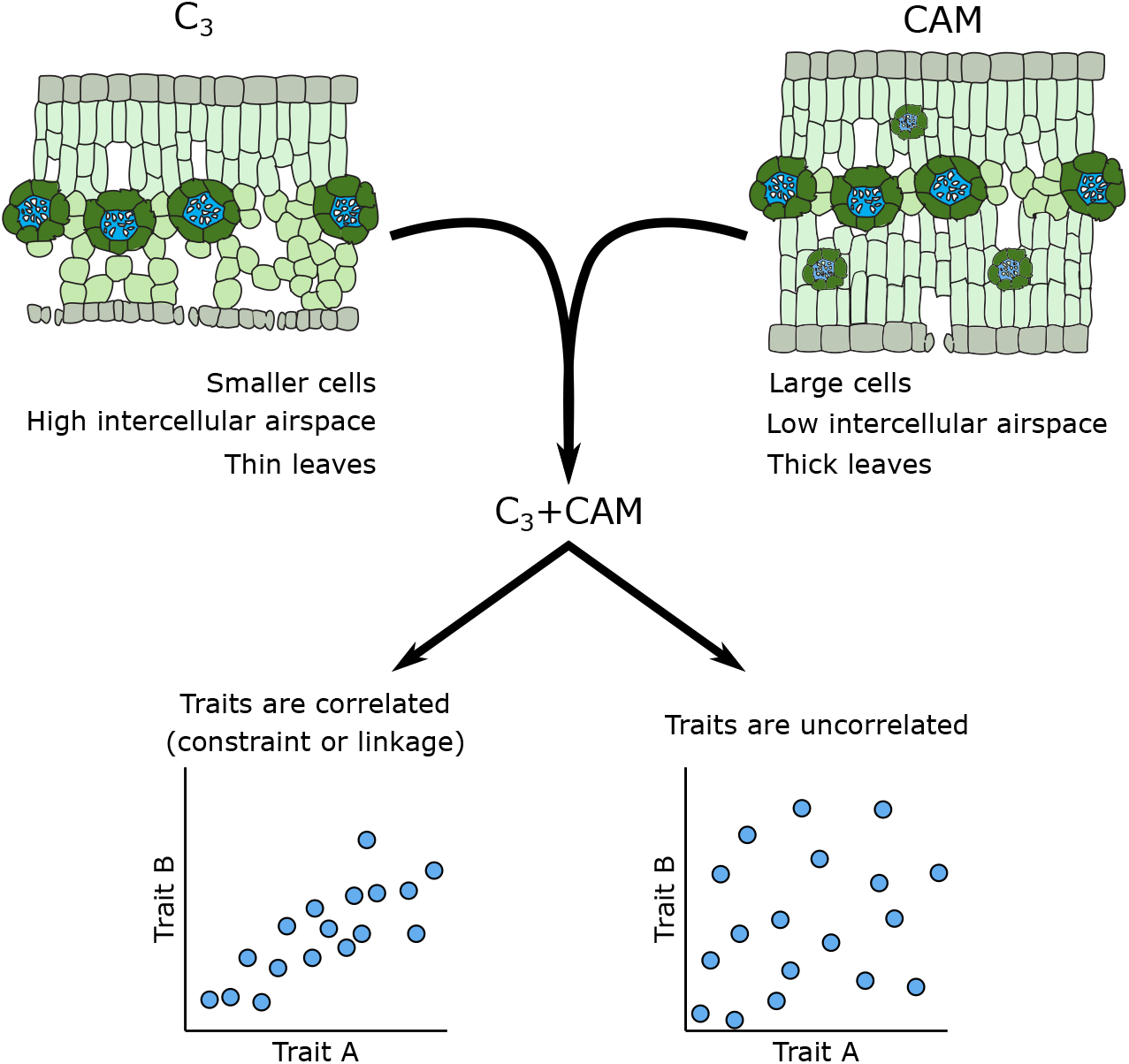
Anatomical traits typically associated with C_3_ and strong CAM plants, showing possible resulting trait associations in a hybrid between a C_3_ and a CAM species.

## Methods

Genotypes of *Yucca gloriosa* were collected from across its range (Virginia to Florida) (Fig. 2) [**Supplementary Table 1**] as ramets, then transplanted to the University of Georgia Department of Plant Biology greenhouses. Plants were grown in 60:40 soil:sand mix with once-weekly watering and fertilizer as needed, and maintained until proper rooting was established and significant new growth was noticeable (minimum 6 months). Beginning in spring 2016, genotypes were randomly assigned to growth chamber experimental runs in sets of four genotypes. For each experimental run, 3-4 clonal replicates for each of the four genotypes were acclimated in the growth chamber for four days prior to manipulation. Growth chambers had 12hr days (beginning at 0700 h), with 30 °C/17 °C day/night temperatures and a relative humidity of 30-40%. Light intensity at leaf level was ~400 μmol m^-2^ s^-2^ and plants were kept well-watered during the acclimation phase. On the first experimental day (“day 1”), gas exchange measurements were collected every two hours for a 24 hour period, beginning one hour after lights turned on, using a LI-6400XT (LI-COR environmental). After day 1, plants were allowed to dry down, soil moisture information was collected on experimental days [**Supplementary Table 2 and Figure 1**], and on day 7 plants were re-measured for gas exchange identically to day 1. At the end of day 7, plants were re-watered, then measured a final time for gas exchange on day 9. Experimental runs were conducted in April, September, and November of 2016, and February, March, April, and August of 2017 on a total of 24 genotypes [**Supplementary Table 2**]. To compare net carbon gain between genotypes, the area under the LI-COR curve per genotype per treatment was calculated using the auc() function in the DescTools (Signorell *et al*. 2019) package in R.

**Figure 2.**
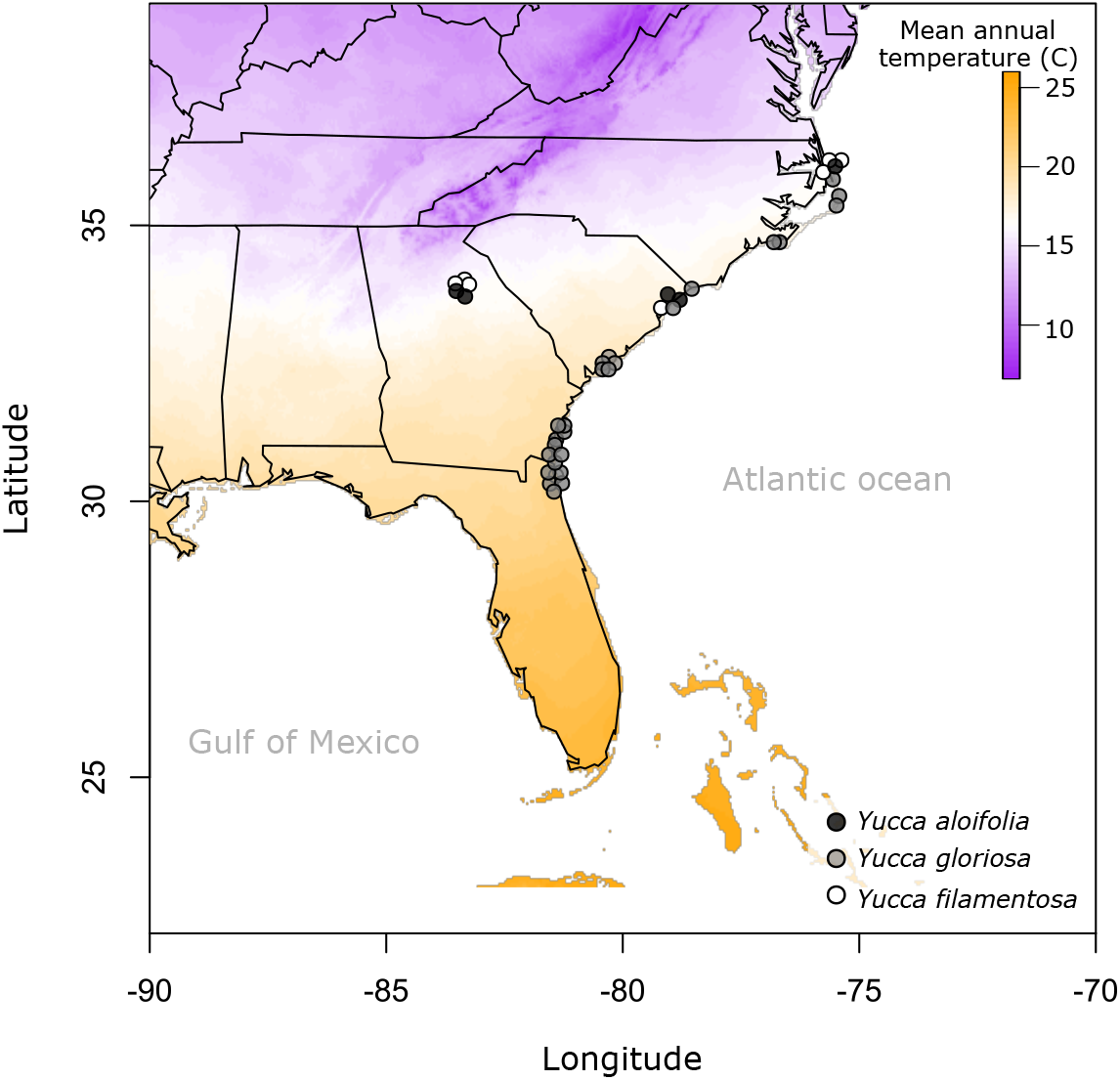
Collection origin of samples used in this study for each of the three *Yucca* species, plotted against mean annual temperature from the Worldclim database. Points are jittered so they do not overlap, see [**Supplementary Table 1**] for full locality information.

Leaf samples for titratable acidity measurements were collected on days 1, 7, and 9 two hours before lights off and two hours before lights turned back on. Leaf punches were taken in triplicate at both time points from all individual plants, then were immediately flash frozen and stored at −80 °C. Leaves were quickly weighed once removed from the freezer and placed into 60 mL of 20% EtOH. Samples were boiled until the volume was reduced by half, at which point the total volume was returned to 60 mL by adding water pH 7.0. Samples were boiled to half volume once more, refilled to 60 mL with water, then allowed to cool to room temperature. The room temperature liquid was cleared of leaf debris and titrated with 0.002 M NaOH to a final pH of 7.0. The umol H+ amount was calculated as (mL of 0.002 M NaOH x 2 mM)/grams of frozen tissue.

Leaf cross sections were collected in April 2018 and April 2019 from clonal replicates of the same genotypes measured for gas exchange, with 2 samples collected per genotype from separate biological replicates when available. Leaves were cut, fixed in formalin, embedded in paraffin, then sectioned at the University of Georgia Veterinary Hospital Histology Lab. Sections were stained with Toluidine Blue and mounted on glass slides. For each of the separate plants sectioned per genotype, 2 images were taken on a Zeiss microscope at 5 and 10x magnification, taking care to avoid imaging edges or damaged sections. Images were analyzed in ImageJ to collect measurements of cell size and intercellular airspace (IAS), as well as leaf thickness. For cell size, the area of five adaxial and abaxial mesophyll cells was measured per image. IAS was measured as a fraction of intercellular air per total cell area and is reported as a percent of mesophyll. Leaf thickness was measured in triplicate across each image analyzed for cell size and IAS. Stomatal density was measured by painting both adaxial and abaxial leaf surfaces with clear nail polish (collected March 2019), then removing with tape and imaging stomatal impressions with a Zeiss microscope. Stomatal measurements were taken from two biological replicates per genotype, when available. Previously collected data on adaxial and abaxial cell sizes from additional genotypes of *Y. gloriosa* was also included (Heyduk, Burrell, *et al*. 2016); while IAS was measured on this previous dataset, due to image quality, we suspect IAS may have been overestimated in the data previously published. IAS was therefore re-analyzed for all data published in 2016. ANOVA or ANCOVAs were performed, as appropriate, to determine the effect of *Y. gloriosa* genotype on phenotypic traits; in the case of CO_2_ uptake and acid accumulation, treatment (watered and drought) was included as a factor.

To compare the hybrid to the parental species, previously published data on *Yucca filamentosa* and *Yucca aloifolia* was included as well (Heyduk, Burrell, *et al*. 2016; Heyduk, Ray, *et al*. 2019). The parental datasets are smaller, in that a total of 5 genotypes (14 individual plants) and 7 genotypes (16 individual plants) were measured for various traits for *Y. aloifolia* and *Y. filamentosa*, respectively. All statistical analyses were conducted in R v. 3.5.1, and raw data is available at www.github.com/kheyduk/Yucca_physiology. We correlated both raw data - that is, individual plant traits - as well as genotypic means using cor.test() in R and adjusting resulting p-values for multiple testing with the Benjamini-Hochberg correction. Correlations were conducted pairwise on all traits, except in cases where one trait was a subset of another (for example, nocturnal CO_2_ total assimilation is a subset of total daily CO_2_ assimilation). Correlations were conducted on individual values and genotypic means of all three species together, then separately for just *Y. gloriosa*. No correlations were tested within the parental species, as the data pulled from earlier work did not have enough replication. For a trait combination to be reported as significant, it had to be significant when correlated both across individuals and across genotypic means; for significant correlations, only the across individual statistics are reported.

## Results

### Gas exchange and titratable acidity

Genotypes of *Yucca gloriosa* varied in their gas exchange patterns over a diel cycle (Fig. 3) [**Supplementary Figure 2**]. The majority of genotypes had some level of C_3_ daytime CO_2_ fixation as well as slight nocturnal CO_2_ assimilation under well-watered conditions (Fig. 3). Under drought stress, overall responses varied. Some genotypes had a nearly total shutdown of gas exchange during the day under drought stress, whereas others maintained non-zero levels. Because plants dried down at slightly variable rates [**Supplementary Table 2 and Figure 1**], we examined the effect of genotype and soil moisture on CO_2_ uptake: well-watered plants had only a slightly significant effect of genotype (F_19,51_=2.16, p<0.05), while drought stressed plants had a significant effect of soil moisture (F_1,48_=15.02, p<0.001). However, nocturnal CO_2_ assimilation (and thus the level of CAM performed) was not significantly related to soil moisture under either well-watered (F_1,51_=0.03, p=0.87) or drought stress (F_1,50_=0.64, p=0.43). Instead, a clear GxE signal was observed via an ANOVA (Type III) assessing interaction of genotype and treatment (well-watered vs. drought) on nocturnal CO_2_ uptake (interaction: F_21,131_=2.36, p<0.01, excluding re-watered measurements) [**Supplementary Table 3**]. At night, certain genotypes (e.g., 16 and 1AB, Fig. 3) were able to increase CO**2** assimilation under drought stress relative to well-watered conditions. No hybrid genotype fully replicated the levels of nocturnal CO_2_ assimilated by *Y. aloifolia*, and many had the ability to use CAM even when well-watered, indicating *Y. gloriosa* is not strictly facultative CAM but rather weak CAM with drought inducibility. Nocturnal acid accumulation in the hybrid *Y. gloriosa*, like gas exchange, had a significant interaction effect between genotype and treatment (F_23,120_=3.73, p<0.001, excluding re-watered measurements) [**Supplementary Table 3**]. In general, the majority of genotypes had an increase in leaf acidity over the night period, indicative of CAM; most genotypes displayed some level of acid accumulation on all days of the experiment, regardless of water status (Fig. 4).

**Figure 3.**
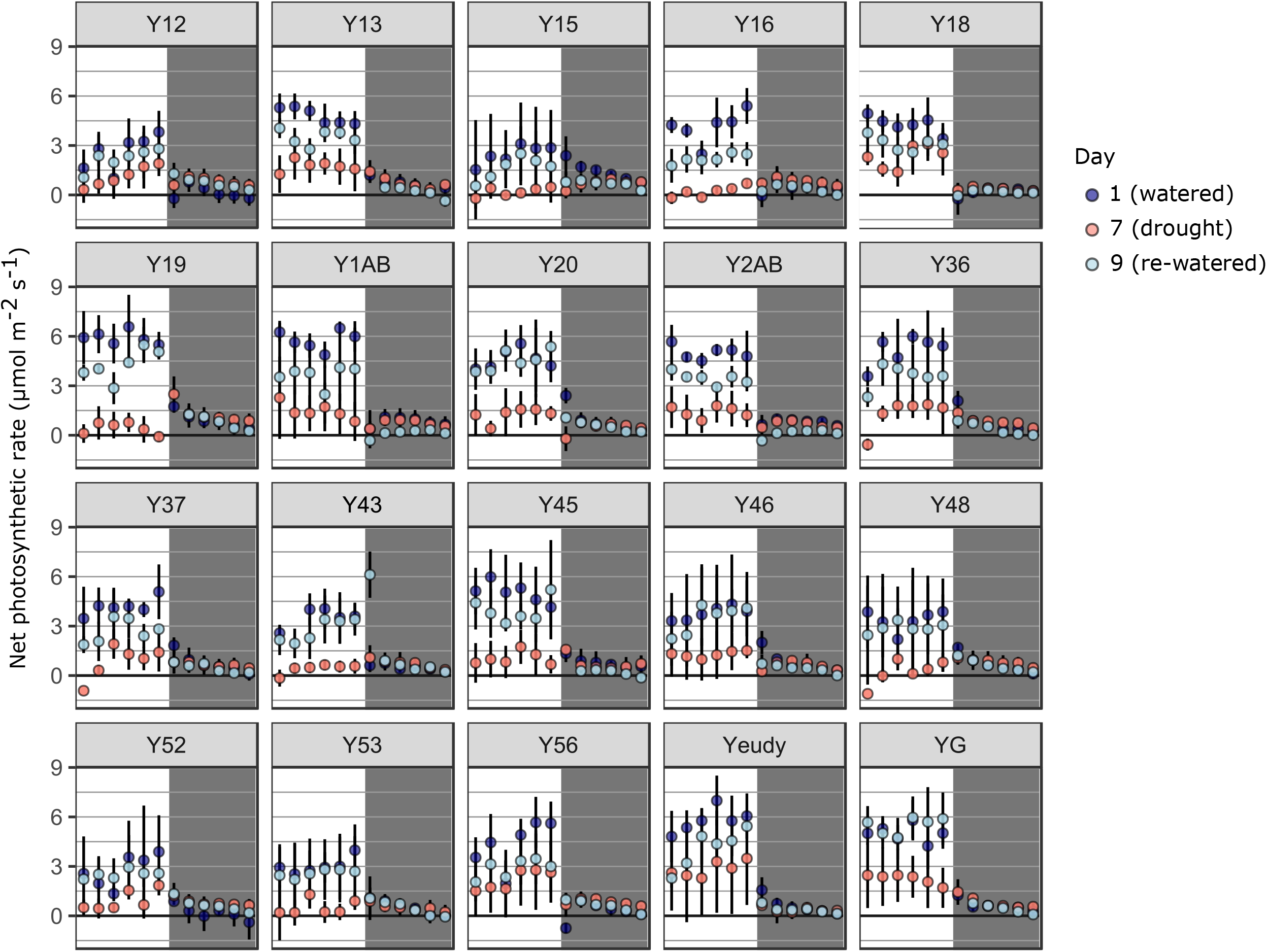
A) Gas exchange of *Y. gloriosa* genotypes measured every two hours over a 24 hour period beginning at 1 hour after lights on (8 a.m.). White and grey backgrounds specify daytime and nighttime measurements, respectively. Mean and standard deviation are shown for days 1 (well-watered), 7 (drought stress), and 9 (re-watered). Four samples were omitted due to potentially inaccurate LiCOR measurements (genotypes 51, 55, 61, and 70).

**Figure 4.**
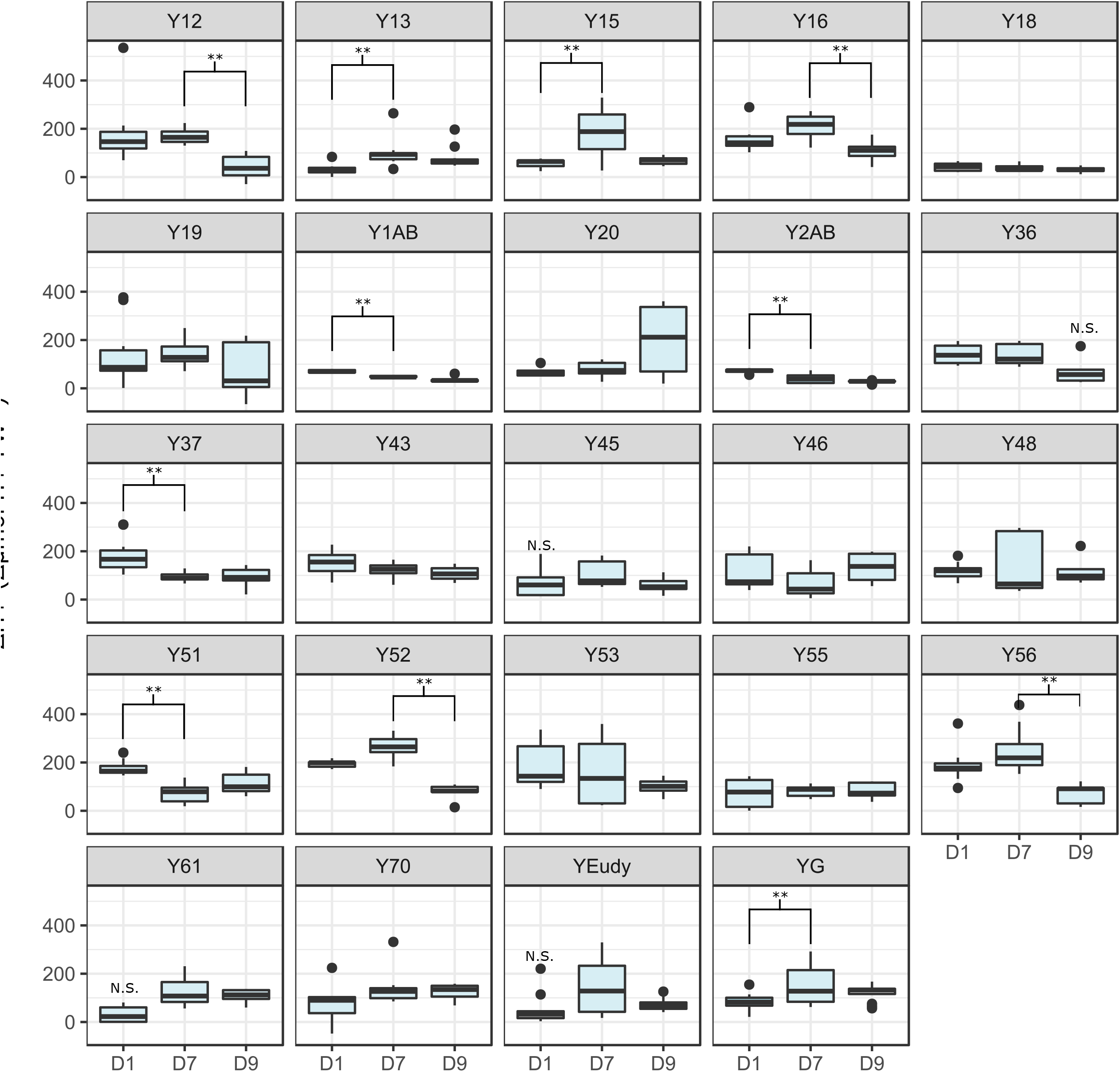
Levels of leaf titratable acidity (AM H+ equivalents - PM H+ equivalents to pH 7.0) across well-watered (D1), drought (D7), and re-watered (D9) time points in 24 genotypes of *Yucca gloriosa*. If any given value is not significantly different from 0 (no acid accumulation), N.S. is shown above the box. Significant difference between time points within a genotype are indicated by brackets above the boxes.

### Inter- and intra-specific response to drought

When compared to the parental genotypes for which gas exchange measurements are available, many *Y. gloriosa* genotypes had some of the highest net CO_2_ assimilation (as calculated by the area under the gas exchange curves) during both well-watered and drought conditions (Fig. 5A). However, nighttime net CO_2_ assimilation was intermediate in *Y. gloriosa* compared to parental species, and tended toward the C_3_ parent *Y. filamentosa* (Fig. 5B). When drought stressed, *Y. filamentosa* genotypes show a decrease in overall CO_2_ assimilation (Fig. 5B) under drought stress. *Yucca aloifolia* showed on average a decrease in nighttime CO_2_ assimilation under drought stress (Fig. 5AB), concordant with previous findings (Heyduk, Burrell, *et al*. 2016). *Yucca gloriosa* genotypes varied in their gas exchange drought response; certain genotypes increased the amount of CO_2_ acquired at night, whereas others, like *Y. aloifolia*, decreased net nighttime CO_2_ acquisition.

**Figure 5.**
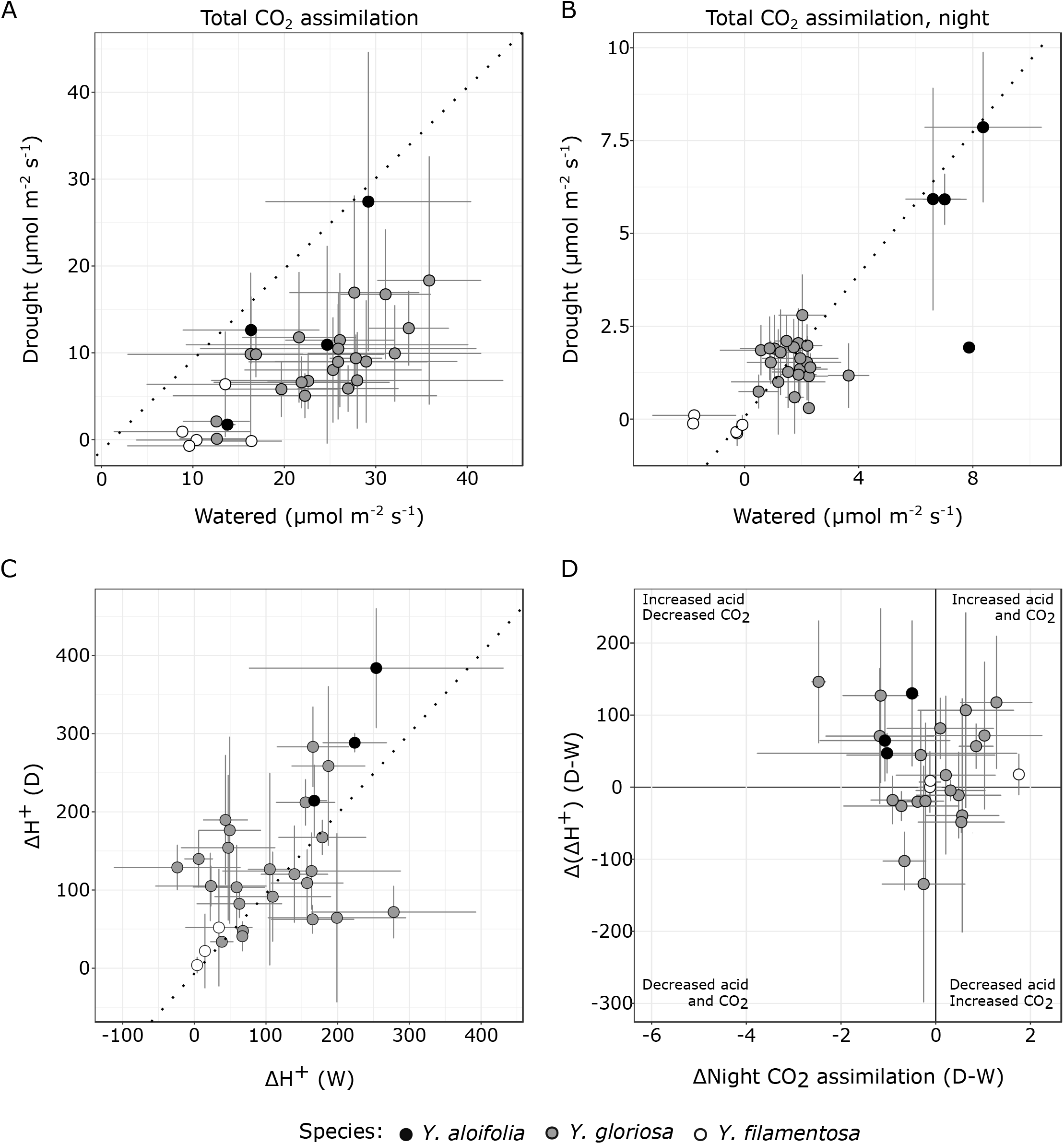
Physiological effect of drought stress on genotypic means (and standard deviation) across all three species. A) The estimated total CO_2_ assimilation, based on the area under the LiCOR curves across the entire diel cycle, for both hybrid and parental genotypes under well-watered and drought-stressed conditions. B) The estimated total CO_2_ assimilation based on the area under the LiCOR curves at night only, for both hybrid and parental genotypes under well-watered and drought-stressed conditions. C) Leaf acid accumulation under well-watered conditions (W) versus drought stressed conditions (D), with genotypic mean and one standard deviation. Dashed line indicates equal values under both conditions; genotypes that fall above or below indicate greater or lower amounts of acid, respectively, were accumulated under drought stress than under well-watered. D) Comparison of the change in nighttime CO_2_ assimilation induced by drought stress (xaxis) to the change in leaf acid accumulation induced by drought (yaxis). Quadrants are labeled with the phenotype observed.

Drought-induced response in leaf acid accumulation varied across hybrid genotypes as well and spanned a larger phenotypic space than either parent. While *Y. filamentosa* never accumulated significantly levels of leaf acid (well-watered: t_5_=1.71, p=0.07; drought stressed: t_5_=1.40, p=0.11), *Y. aloifolia* had appreciable levels of acid accumulation over the night period under well-watered conditions and increased the amount of acid accumulated under drought (Fig. 5B). *Yucca gloriosa* genotypes spanned the range from low levels of acid accumulation to CAM-like levels under well-watered conditions, and genotypes varied in their ability to increase the amount of acid stored under drought. A few genotypes responded to drought with significant and positive increases in leaf acidity on day 7 relative to day 1 (e.g., Y13 and YG). Genotype Y18 is a notable exception in its lack of acid accumulation and lack of response to drought stress, which corresponds to its negligible rates of CO_2_ assimilation during the dark period (Fig. 3). Genotypes that had high levels of acid accumulation under well-watered conditions tended to decrease acid under drought, while those that had lower level well-watered acid accumulation tended to increase the amount of acid stored in leaves under drought stress (Fig. 5B).

Because CAM can be defined by both acid accumulation and nighttime CO_2_ assimilation, comparing the response of genotypes through both phenotypes can indicate the mode of CAM employed under drought stress. For example, *Y. aloifolia* reduces the amount of CO_2_ assimilated at night, but typically increases leaf acid accumulation (Fig. 5D), indicating more reliance on recycling CO_2_ while drought stressed. Some genotypes of *Y. gloriosa* decreased reliance on atmospheric CO**2** and increase acid accumulation with drought stress (upper left quadrant, Fig. 5D). Others responded to drought by increasing both nighttime CO_2_ assimilation and leaf acid accumulation (upper right quadrant, Fig. 5D). A few genotypes were negatively impacted by drought in that they reduced both leaf acid accumulation and nighttime CO_2_ uptake, such that stress appears to have diminished their CAM capacity (lower left quadrant, Fig. 5D). Finally, a few genotypes appear to increase the amount of nighttime CO_2_ assimilation but *decrease* the level of acid stored in the leaves (lower right quadrant, Fig. 5D); however in many of these latter cases the error bars overlap zero, and therefore we cannot reject the expectation that nighttime CO_2_ uptake is coupled with acid accumulation in these genotypes. Regardless, the general diversity of drought responses in the hybrid *Y. gloriosa* is clear.

### Leaf anatomy

All anatomical traits were significantly different between species, based on ANOVA (Benjamini-Hochberg corrected p-values) [**Supplementary Table 4**], with the exception of abaxial stomatal density. Within *Y. gloriosa*, the only anatomical traits significantly different between genotypes were IAS (F_25,21_=3.36, p<0.001) and mean stomatal density (averaged abaxial and adaxial values) (F_4,13_=3.73, p<0.01) [**Supplementary Table 3**]. As with physiological traits, anatomical differences between *Y. aloifolia* and *Y. filamentosa* were stark, while the hybrid largely filled the phenotypic space between. Cell sizes on both adaxial and abaxial sides of the leaf were larger in *Y. aloifolia* than in either of the other two species (Fig. 5A). Stomatal density, conversely, was lowest on average in *Y. aloifolia* and greatest in *Y. filamentosa* (Fig. 6B). Cell sizes and stomatal densities on adaxial and abaxial sides of the leaf were highly positively correlated (Fig. 6AB) across all individuals (cell size: t_67_=25.19, R^2^= 0.90, p<0.001; stomata: t_46_=7.12, R^2^=0.52, p<0.001).

**Figure 6.**
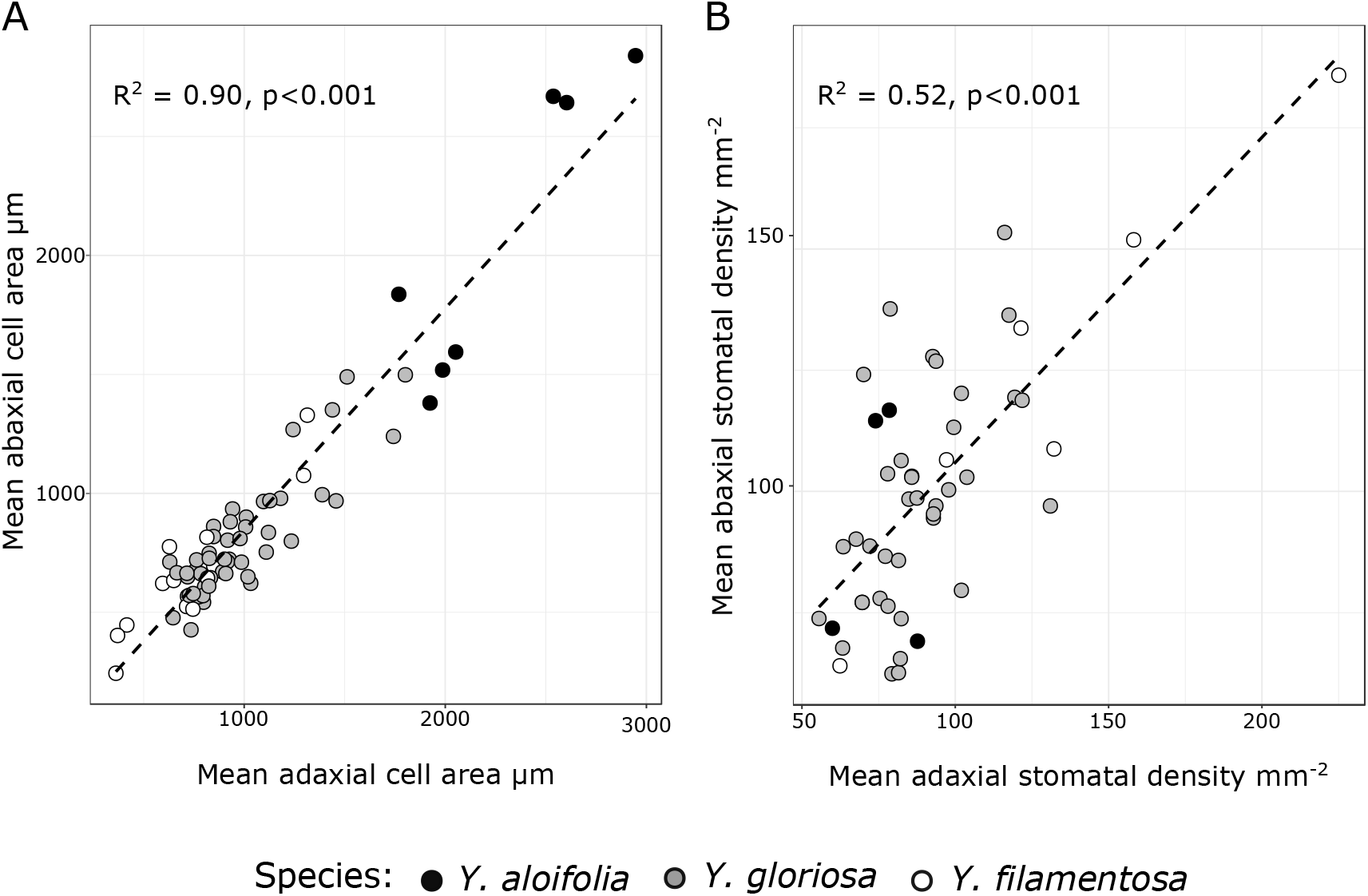
Mean values of cell sizes (A) and stomatal densities (B) on adaxial and abaxial sides of the leaf per individual plant. In both cases, the dashed line is the regression line from the lm() function in R. R^2^ and p-value are reported based on correlation tests in R.

Few anatomical traits could predict physiological traits across species (Fig. 7) [**Supplementary Table 5 and Figure 3**]. Total CO_2_ assimilation across the whole day under both water and drought stress was correlated to IAS, albeit with a relatively low R^2^ in both cases (Fig. 7A). The amount of nocturnal CO_2_ assimilated under both watered and drought conditions had multiple trait correlations to both other physiological traits as well as leaf anatomy. Nocturnal CO_2_ uptake was correlated to IAS, levels of acid accumulation (both under watered and drought conditions), the maximum amount of acid held within a leaf at any time point, the mean cell size, and leaf thickness (Fig. 7B-G). The amount of leaf acids accumulated under drought stress was correlated to total nocturnal CO_2_ assimilation under drought stress (R^2^=0.14, p<0.01). Within *Y. gloriosa*, nearly all the trait correlations were not significant [**Supplementary Table 6**]. The only significant correlations for traits in *Y. gloriosa* were between mean cell size and leaf thickness (R^2^=0.57, p<0.001) and between total CO_2_ assimilation under water and drought conditions (R^2^=0.24, p<0.001).

**Figure 7.**
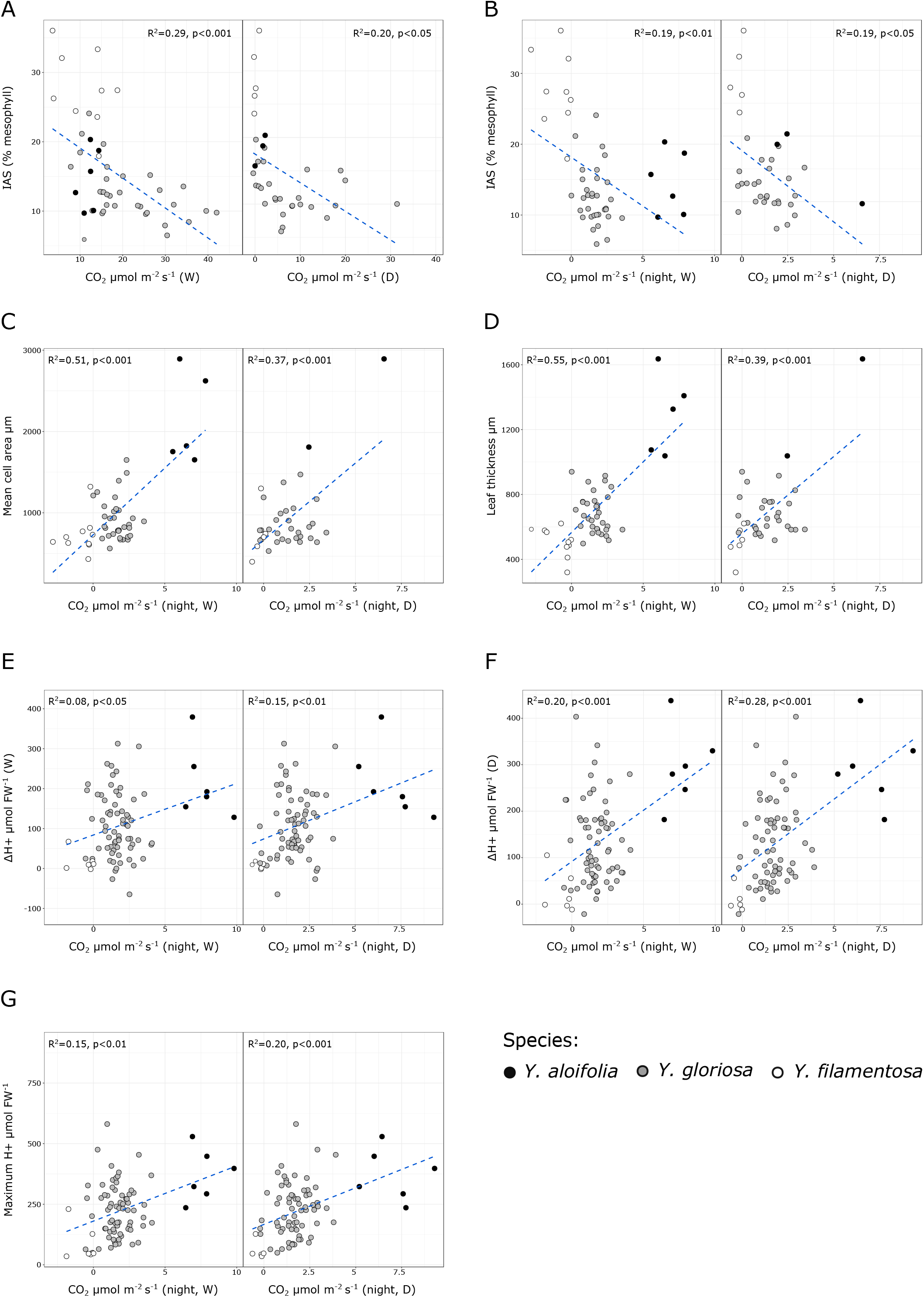
Scatterplots and regression lines with R^2^ and p-values for a subset of traits [**Supplementary Figure 3**]. Individual data per plant are plotted, and correlations are shown for individual plant data, rather than genotypic means [**Supplementary Table 5**]. A) Total CO_2_ assimilation under watered and drought plotted against intercellular airspace (IAS). B-G) Nocturnal CO_2_ assimilation under watered and drought plotted against IAS (B), mean cell area (C), leaf thickness (D), leaf acid accumulation under watered (E) and drought (F) treatments, and against the maximum amount of acid present at any time point (G).

## Discussion

Detailed physiological and anatomical measurements in *Y. gloriosa* have revealed among-genotype variation in CAM phenotypes, and that anatomical and physiological traits show a lack of correlation within *Y. gloriosa*. Under drought stress, the levels of daytime CO_2_ assimilation were largely driven by environment - that is, soil moisture content - whereas nocturnal CO_2_ assimilation rates and acid accumulation were influenced by a combination of genotype and environmental effects. Our results reveal a continuum of photosynthetic traits across *Y. gloriosa* genotypes, including variation in drought response. Anatomical measurements were largely not predictive of physiological traits within *Y. gloriosa*. In contrast, cell size, IAS, and leaf thickness were predictive of nocturnal CO_2_ uptake in cross species comparisons. These observations suggest that anatomical characteristics can be decoupled from photosynthetic physiology of CAM within the homoploid hybrid species *Y. gloriosa*.

### Intra- and interspecific variation in anatomy and physiology

The few studies that have linked leaf anatomy to CAM photosynthetic capacity have provided often contrasting results on how important various anatomical traits are for CAM. In a study that compared phylogenetically unrelated strong CAM and C_3_+CAM species, cell size was found to be unrelated to CAM (Nelson and Sage 2008). In contrast, a study of *Clusia* species that ranged from C_3_ to CAM with intermediates showed palisade mesophyll cell size was significantly correlated to the proportion of CO_2_ uptake at night (Zambrano *et al*. 2014). Across the three *Yucca* species examined here, cell size (area) was related to nocturnal CO_2_ uptake under both watered and drought conditions, although such a relationship did not exist at the intraspecific level within *Y. gloriosa*. All studies use cell size as a proxy for vacuolar size, which in theory would limit the storage capacity of malate. It is possible that vacuolar size is not linearly related to cell size (though see Chan and Marshall 2014), and that inconsistent results on the importance of cell size between studies is related to using anatomical proxies for the true trait of interest. Alternatively, and probably more likely, studies that control for phylogenetic distance, such as this one and those conducted across *Clusia* species, reduce noise introduced by sampling across evolutionarily distant lineages and may provide a more accurate assessment of anatomical importance.

In addition to cell size, intercellular air space (IAS) is often cited as a critical trait for CAM, though whether it evolves as a byproduct of tight cell packing (Maxwell *et al*. 1997) or as a way to reduce CO_2_ remains unclear (Borland *et al*. 2018). IAS was strongly correlated to strength of CAM when measured across unrelated CAM and C_3_+CAM species (Nelson and Sage 2008), but had little role in determining strength of CAM when assessed within the genus *Clusia* (Zambrano et al. 2014). IAS was correlated to nocturnal CO_2_ assimilation when tested across all three species of *Yucca*, but was not correlated to leaf acid accumulation, and showed no relationship to *any* other traits within *Y. gloriosa*. That IAS is not predictive of physiology in *Y. gloriosa* - neither nocturnal CO_2_ uptake or the amount of leaf acids that accumulate - is surprising, given that contrasts between C_3_ to CAM species have repeatedly shown the latter have significantly reduced IAS (Heyduk, McKain, *et al*. 2016; Heyduk, Burrell, *et al*. 2016; Males 2018). Together, the IAS trends across and within *Yucca* species shows that while IAS may be required for constitutive CAM, there exists a large intermediate space where IAS predicts very little about photosynthetic functionality.

For many anatomical and physiological traits, *Y. gloriosa* has intermediate values compared to the two parental species and often occupies a much broader range of trait values. It is possible that our limited sampling of the parental species, drawn from previous work, reduces our ability to accurately assess trait space in *Y. aloifolia* and *Y. filamentosa*. However, multiple genotypes were sampled across the ranges of both parental species, and thus the greater variation found within the hybrid is likely due to genomic mixing, rather than sampling bias. While trait values in *Y. gloriosa* were typically intermediate, one notable exception was the transgressive values of total CO_2_ assimilation under both watered and drought stressed conditions. Due to *Y. gloriosa*’s nearly C_3_-level of daytime CO_2_ fixation coupled with the ability to use low level CAM, total CO_2_ uptake rates far exceed that of either parent, at least in certain genotypes. Such a mixed photosynthetic strategy may be particularly valuable on the coastal dunes that *Y. gloriosa* is restricted to, as although rainfall in the southeastern U.S. is not particularly limiting, any water that does fall likely percolates through the sandy substrate quickly.

Despite the potentially novel phenotypes that *Y. gloriosa* exhibits relative to its parental species, they are unlikely to underlie the speciation of the hybrid from its progenitors. All three *Yucca* species are found across the southeastern coast, although only *Y. aloifolia* and *Y. gloriosa* grow with any frequency on the coastal dunes. *Y. filamentosa* is typically further away from the ocean in the coastal scrub, though can be found in exposed sand near brackish inlets (K Heyduk, unpubl. res.). Homoploid hybrid species can be formed and maintained either through chromosomal structural rearrangements that form a reproductive barrier between the new species and its progenitors, or via ecological differentiation, whereby the new combination of traits in the hybrid allows for habitation of a novel niche relative to the parental species (Gross and Rieseberg 2005). As the habitats of the *Yucca* species studied here largely overlap, particularly *Y. gloriosa* and *Y. aloifolia*, the latter at first seems unlikely, despite *Y. gloriosa* being clearly distinct in total CO_2_ assimilation rates. However, flowering time of the three species is markedly different: *Y. filamentosa* typically flowers earliest, in late May and June, followed by *Y. aloifolia. Yucca gloriosa* has been noted to flower at the end of the summer and into autumn (Trelease 1902); whether the later flowering time was selected for in order to reduce backcrossing, or was instead a byproduct of the initial hybridization events, remains unknown. Additionally, other biotic interactions (e.g., below-ground mutualisms or pollinators) or microhabitat variation are largely untested as potential drivers of *Yucca* speciation (but see Rentsch and Leebens-Mack 2014). Chromosomal structural rearrangements may explain an apparent lack of backcrossed individuals, but we do not currently have the genomic data to test this hypothesis.

### Abiotic stress regulation of CAM

Genotypes of *Y. gloriosa* used variable levels of CAM, and differentially up-regulated CAM under drought stress. The differential drought response was a result of two separate axes of the CAM phenotype: both leaf acid accumulation and nocturnal CO_2_ uptake varied by environment, and could do so in a de-coupled manner (Fig. 5). That is, certain genotypes increased the amount of leaf acids accumulated based not on increasing atmospheric CO_2_ uptake but instead by presumably re-fixing respired CO_2_. Such a response indicates that much of the required enzymes are present and regulated correctly, but that stomatal aperture responded negatively to drought at night. Reducing net CO_2_ uptake but increasing leaf acid accumulation is the typical response of *Y. aloifolia* to drought stress as well. In general, the response to drought stress in *Y. gloriosa* was transgressive relative to parental phenotypes, in that genotypes of *Y. gloriosa* were able to respond to drought stress in ways that neither parent could. For example, certain genotypes could *increase* both CO_2_ uptake and leaf acid accumulation under drought stress - this response was not seen in any of the parental genotypes measured here. Other genotypes occupied a part of trait space where nocturnal CO_2_ uptake increased under drought stress, but leaf acids decreased (Fig. 5D). How incoming CO_2_ is processed in these genotypes remains unclear and warrants additional exploration in these genotypes, especially through metabolomic and genomic analyses to help pinpoint alternative pathways.

The segregation of CAM drought response in *Y. gloriosa* also presents an ideal system with which to interrogate the molecular components of drought response in facultative CAM species. CAM has been touted as a potential trait for increasing food and biofuel crop drought tolerance through bioengineering (Borland *et al*. 2014, 2015), and early efforts to transform C_3_ species to CAM were instrumental in generating an abundance of genomics data for CAM species. Yet fully committing a C_3_ plant to CAM may result in costs that outweigh any gains in drought tolerance; larger leaves and cells will require more energy and time to produce, and constitutive CAM usage is not ideal when drought may be intermittent. Instead, drought tolerance engineering efforts should look to facultative CAM or C_3_+CAM, as in *Y. gloriosa*, which outperforms its parental species in terms of total CO_2_ uptake and may result in faster biomass gains, though this remains to be tested. The natural variation for CAM induction and up-regulation in *Y. gloriosa*, as well as the uncoupling of various CAM traits, including anatomy, acid accumulation, and CO_2_ uptake, make *Y. gloriosa* ideal for investigating the molecular basis of particular CAM traits and their regulation via abiotic signaling.

### Implications for the evolution of CAM

While *Y. gloriosa* is a hybrid and therefore represents a somewhat atypical avenue for trait evolution, it still allows us a glimpse into how a trait like CAM might be assembled. The homoploid nature of *Y. gloriosa* means that the genomic content of the two parental species is not highly divergent, and that the mix of traits found in *Y. gloriosa* genotypes are not a result of a highly perturbed genome but more like what may be expected of an intraspecific cross between phenotypically divergent parents. The mixture of traits within *Y. gloriosa* allows us to speculate on the genomic architecture underlying the CAM phenotype. It seems unlikely that many of the traits are genetically linked - that is, the few relationships between traits within *Y. gloriosa* means the underlying genes are dispersed in physical genomic location and that they are not necessarily expressed in or regulated by similar pathways. For example, there is nothing in the genome of *Y. gloriosa* that requires large cells to develop low IAS (or vice versa), or that CAM activity is in any way linked genetically to leaf thickness. The variation in and lack of association between traits in *Y. gloriosa* also implies, unsurprisingly, that the CAM phenotype is highly quantitative, and that recombination can break up many of the underlying traits. The genetic architecture of CAM does not fully explain why such a mix of traits has remained in *Y. gloriosa*. The frequent dry down of sandy coastal dunes may be promoting the maintenance of intermediate traits as a way of mediating a highly variable environment. Alternatively, *Y. gloriosa* is not a particularly rare species in its native habitat, but its populations are small and relatively isolated. In such small populations, selection has a weaker effect than drift, which can lead to less advantageous combinations of traits persisting in a species. Additional research using common gardens could facilitate our understanding of whether intermediate traits like those found in *Y. gloriosa* can confer a fitness advantage in some circumstances.

The variation and lack of correlation between traits underlying the CAM phenotype in *Y. gloriosa* also give insight into how CAM is assembled over evolutionary time. While certain traits appear fixed when we examine strong C_3_ and CAM species, intermediate species are important for understanding the processes that may have led to trait fixation and correlation across traits. After all, selection acts not on the species level, but on individual, and indeed there is a broad phenotypic space within *Y. gloriosa* for the traits examined here that selection could act upon. That selection seems to recurrently end up on a particular anatomical phenotype in CAM species - i.e., larger cells, thicker leaves - despite no genetic constraint for such a correlation suggests there is an optimal combination of traits for CAM efficiency. The pattern of convergent evolution of trait combinations, paired with intermediate species showing highly variable trait combinations, implies a funnel shape to the evolutionary trajectory of CAM. Species can use a degree of CAM without committing to any particular leaf anatomy, meaning that initial transitions to using C_3_+CAM following broad and varied routes. This is in contrast to the evolution of C_4_ photosynthesis, which, like CAM, requires specific anatomical characteristics. In C_4_ lineages, anatomical changes occur prior to the evolution of C_4_ biochemistry (McKown and Dengler 2007; Lundgren *et al*. 2019); in some cases, like the PACMAD clade of grasses, these anatomical changes happen early enough in evolutionary time that they are thought to have facilitated repeated origins of C_4_ (Christin *et al*. 2013). In contrast, “weak” CAM or C_3_+CAM has no anatomical constraints in *Yucca*. There is, however, an upper bound where further investment in CO_2_ fixation by the CAM pathway requires dedicated anatomy, though the exact threshold of that transition point remains unclear. The funnel shape to the evolution of CAM, whereby no anatomical constraints impact low levels of CAM function, means that ordering of events on the evolutionary trajectory from C_3_ to CAM will be exceedingly difficult, as lineages can take various routes through the intermediate zone.

While the lack of correlated traits in an intermediate C_3_+CAM hybrid species has implications for broader questions on the evolution of CAM, future work can elaborate upon the patterns seen here and help assess how generalizable these results are. Sampling of parental genotypes and overall range was limited, and thus there may exist greater variation among traits in the parental C_3_ and CAM species as well. Indeed, most studies that examine the correlation of anatomy to photosynthetic physiology do not sample intraspecific variation, and therefore it remains a largely unexplored area of CAM. Growth conditions used in this study were based on previous work in the *Yucca* system, but modulating those conditions may reveal deeper levels of variation across environmental gradients. Finally, the *Yucca* hybrid system is a single example of intermediacy between C_3_ and CAM, and other C_3_+CAM species should continue to be examined via detailed physiology and anatomy to advance fundamental understanding of how CAM evolves. Investigations within and between species exhibiting a mix of CAM, C_3_, and intermediate species will continue to provide insights into whether the decoupling of CAM traits we observe in a hybrid species holds more generally.

### Conclusions

Comparisons between C_3_ and CAM species have suggested suites of traits are correlated to maximize the efficiency of each photosynthetic pathway: CAM species have large cells for storing malate, and the cells are often packed together densely in large, thick leaves to minimize CO_2_ leakage back into the atmosphere; C_3_ plants have large amounts of airspace between significantly smaller cells to facilitate the diffusion of CO_2_ to the sites of Rubisco carboxylation. These trends have been seen repeatedly in independent CAM lineages, but few studies have examined intermediate C_3_+CAM plants, and even fewer have assessed intraspecific variation for trait correlations. The C_3_+CAM hybrid *Y. gloriosa* examined here not only has a greater range of traits than either of its parental species, but it also lacks many of the trait correlations commonly associated with the ability to use CAM. Indeed, not a single leaf anatomical trait could predict the amount of CO_2_ acquired via CAM in the hybrid species. The lack of variation within the intermediate *Y. gloriosa* suggests that the evolutionary trajectory to CAM from C_3_ passes through a stage where many combinations of anatomical and photosynthetic physiology traits can coexist in a single plant. Furthermore, in *Yucca* at least, anatomical and physiological traits are not genetically linked, and supports existing hypotheses that suites of leaf traits found repeatedly in CAM species have been selected for in order to maximize photosynthetic efficiency.

## Acknowledgements

The authors would like to thank the state parks of Georgia, Florida, South Carolina, North Carolina, and Virginia, as well as the National Park Service, for their assistance in collection and permitting. Special thanks to Nida Moledina, Dan Tepstov, Grace Manning, Rushi Patel, and Charmi Patel for measuring thousands of titration samples, the greenhouse staff at UGA (particularly Mike Boyd and Greg Cousins) for granting us growth chamber access, and Rick Field and Amanda Cummings for assistance in collecting anatomical samples.

## References

Borland AM, Hartwell J, Weston DJ, et al. 2014. Engineering crassulacean acid metabolism to improve water-use efficiency. Trends in plant science 19: 327–338.

Borland AM, Leverett A, Hurtado-Castano N, Hu R, Yang X. 2018. Functional Anatomical Traits of the Photosynthetic Organs of Plants with Crassulacean Acid Metabolism In: Adams WW III, Terashima I, eds. The Leaf: A Platform for Performing Photosynthesis. Cham: Springer International Publishing, 281–305.

Borland AM, Wullschleger SD, Weston DJ, et al. 2015. Climate-resilient agroforestry: physiological responses to climate change and engineering of crassulacean acid metabolism (CAM) as a mitigation strategy. Plant, cell & environment 38: 1833–1849.

Bradshaw HD, Schemske DW. 2003. Allele substitution at a flower colour locus produces a pollinator shift in monkeyflowers. Nature 426: 176–178.

Chan Y-HM, Marshall WF. 2014. Organelle size scaling of the budding yeast vacuole is tuned by membrane trafficking rates. Biophysical journal 106: 1986–1996.

Christin P-A, Osborne CP, Chatelet DS, et al. 2013. Anatomical enablers and the evolution of C4 photosynthesis in grasses. Proceedings of the National Academy of Sciences of the United States of America 110: 1381–1386.

Edwards EJ. 2019. Evolutionary trajectories, accessibility, and other metaphors: the case of C4 and CAM photosynthesis. The New phytologist.

Edwards EJ, Ogburn RM. 2012. Angiosperm Responses to a Low-CO 2 World: CAM and C 4 Photosynthesis as Parallel Evolutionary Trajectories. International journal of plant sciences 173: 724–733.

Goolsby EW, Moore AJ, Hancock LP, De Vos JM, Edwards EJ. 2018. Molecular evolution of key metabolic genes during transitions to C4 and CAM photosynthesis. American journal of botany 105: 602–613.

Gross BL, Rieseberg LH. 2005. The ecological genetics of homoploid hybrid speciation. The Journal of heredity 96: 241–252.

Hartsock TL, Nobel PS. 1976. Watering converts a CAM plant to daytime CO2 uptake. Nature 262: 574–576.

Hatch MD. 1987. C4 photosynthesis: a unique blend of modified biochemistry, anatomy and ultrastructure. Biochimica et Biophysica Acta (BBA) - Reviews on Bioenergetics 895: 81–106.

Herrera A. 2009. Crassulacean acid metabolism and fitness under water deficit stress: if not for carbon gain, what is facultative CAM good for? Annals of botany 103: 645–653.

Heyduk K, Burrell N, Lalani F, Leebens-Mack J. 2016. Gas exchange and leaf anatomy of a C3-CAM hybrid, Yucca gloriosa (Asparagaceae). Journal of experimental botany 67: 1369–1379.

Heyduk K, Hwang M, Albert V, et al. 2018. Altered Gene Regulatory Networks Are Associated With the Transition From C3 to Crassulacean Acid Metabolism in Erycina (Oncidiinae: Orchidaceae). Frontiers in plant science 9: 2000.

Heyduk K, McKain MR, Lalani F, Leebens-Mack J. 2016. Evolution of CAM anatomy predates the origins of Crassulacean acid metabolism in the Agavoideae (Asparagaceae). Molecular phylogenetics and evolution 105: 102–113.

Heyduk K, Moreno-Villena JJ, Gilman IS, Christin P-A, Edwards EJ. 2019. The genetics of convergent evolution: insights from plant photosynthesis. Nature reviews. Genetics 2019.

Heyduk K, Ray JN, Ayyampalayam S, et al. 2019. Shared expression of Crassulacean acid metabolism (CAM) genes predates the origin of CAM in the genus Yucca. Journal of experimental botany.

Hoekstra HE, Hirschmann RJ, Bundey RA, Insel PA, Crossland JP. 2006. A single amino acid mutation contributes to adaptive beach mouse color pattern. Science 313: 101–104.

Lee H, Rustgi S, Kumar N, et al. 2011. Single nucleotide mutation in the barley acetohydroxy acid synthase (AHAS) gene confers resistance to imidazolinone herbicides. Proceedings of the National Academy of Sciences of the United States of America 108: 8909–8913.

Lenski RE, Ofria C, Pennock RT, Adami C. 2003. The evolutionary origin of complex features. Nature 423:139–144.

Lundgren MR, Dunning LT, Olofsson JK, et al. 2019. C4 anatomy can evolve via a single developmental change. Ecology letters 22: 302–312.

Males J. 2018. Concerted anatomical change associated with crassulacean acid metabolism in the Bromeliaceae. Functional plant biology: FPB 45: 681–695.

Maxwell K, Caemmerer S von, Evans JR. 1997. Is a Low Internal Conductance to CO2 Diffusion a Consequence of Succulence in Plants with Crassulacean Acid Metabolism? Australian journal of plant physiology 24: 777.

McKown AD, Dengler NG. 2007. Key innovations in the evolution of Kranz anatomy and C4 vein pattern in Flaveria (Asteraceae). American journal of botany 94: 382–399.

Ming R, VanBuren R, Wai CM, et al. 2015. The pineapple genome and the evolution of CAM photosynthesis. Nature genetics 47: 1435–1442.

Nelson EA, Sage RF. 2008. Functional constraints of CAM leaf anatomy: tight cell packing is associated with increased CAM function across a gradient of CAM expression. Journal of experimental botany 59: 1841–1850.

Nelson EA, Sage TL, Sage RF. 2005. Functional leaf anatomy of plants with crassulacean acid metabolism. Functional plant biology: FPB 32: 409.

Rentsch JD, Leebens-Mack J. 2012. Homoploid hybrid origin of Yucca gloriosa: intersectional hybrid speciation in Yucca (Agavoideae, Asparagaceae). Ecology and evolution 2: 2213–2222.

Rentsch JD, Leebens-Mack J. 2014. Yucca aloifolia (Asparagaceae) opts out of an obligate pollination mutualism. American journal of botany 101: 2062–2067.

Sage RF, Khoshravesh R, Sage TL. 2014. From proto-Kranz to C4 Kranz: building the bridge to C4 photosynthesis. Journal of experimental botany 65: 3341–3356.

Signorell A. 2019. DescTools: Tools for Descriptive Statistics.

Silvera K, Santiago LS, Winter K. 2005. Distribution of crassulacean acid metabolism in orchids of Panama: evidence of selection for weak and strong modes. Functional plant biology: FPB 32: 397.

Silvera K, Winter K, Rodriguez BL, Albion RL, Cushman JC. 2014. Multiple isoforms of phosphoenolpyruvate carboxylase in the Orchidaceae (subtribe Oncidiinae): implications for the evolution of crassulacean acid metabolism. Journal of experimental botany 65: 3623–3636.

Trelease W. 1902. The yucceae. Missouri Botanical Garden Annual Report 1902: 27–133.

Wai CM, Weise SE, Ozersky P, Mockler TC, Michael TP, VanBuren R. 2019. Time of day and network reprogramming during drought induced CAM photosynthesis in Sedum album. PLoS genetics 15:e1008209.

Weber JN, Peterson BK, Hoekstra HE. 2013. Discrete genetic modules are responsible for complex burrow evolution in Peromyscus mice. Nature 493: 402–405.

Winter K. 2019. Ecophysiology of constitutive and facultative CAM photosynthesis. Journal of experimental botany.

Winter K, Garcia M, Holtum JAM. 2008. On the nature of facultative and constitutive CAM: environmental and developmental control of CAM expression during early growth of Clusia, Kalanchöe, and Opuntia. Journal of experimental botany 59: 1829–1840.

Yang X, Hu R, Yin H, et al. 2017. The Kalanchoë genome provides insights into convergent evolution and building blocks of crassulacean acid metabolism. Nature communications 8: 1899.

Yin H, Guo H-B, Weston DJ, et al. 2018. Diel rewiring and positive selection of ancient plant proteins enabled evolution of CAM photosynthesis in Agave. BMC genomics 19: 588.

Yuan Y-W, Sagawa JM, Young RC, Christensen BJ, Bradshaw HD Jr. 2013. Genetic dissection of a major anthocyanin QTL contributing to pollinator-mediated reproductive isolation between sister species of Mimulus. Genetics 194: 255–263.

Zambrano VAB, Lawson T, Olmos E, Fernández-García N, Borland AM. 2014. Leaf anatomical traits which accommodate the facultative engagement of crassulacean acid metabolism in tropical trees of the genus Clusia. Journal of experimental botany 65: 3513–3523.

