## Supplementary Figure 1 for "Leaf anatomy is not correlated to CAM function in a C_3_+CAM hybrid species, *Yucca gloriosa*"

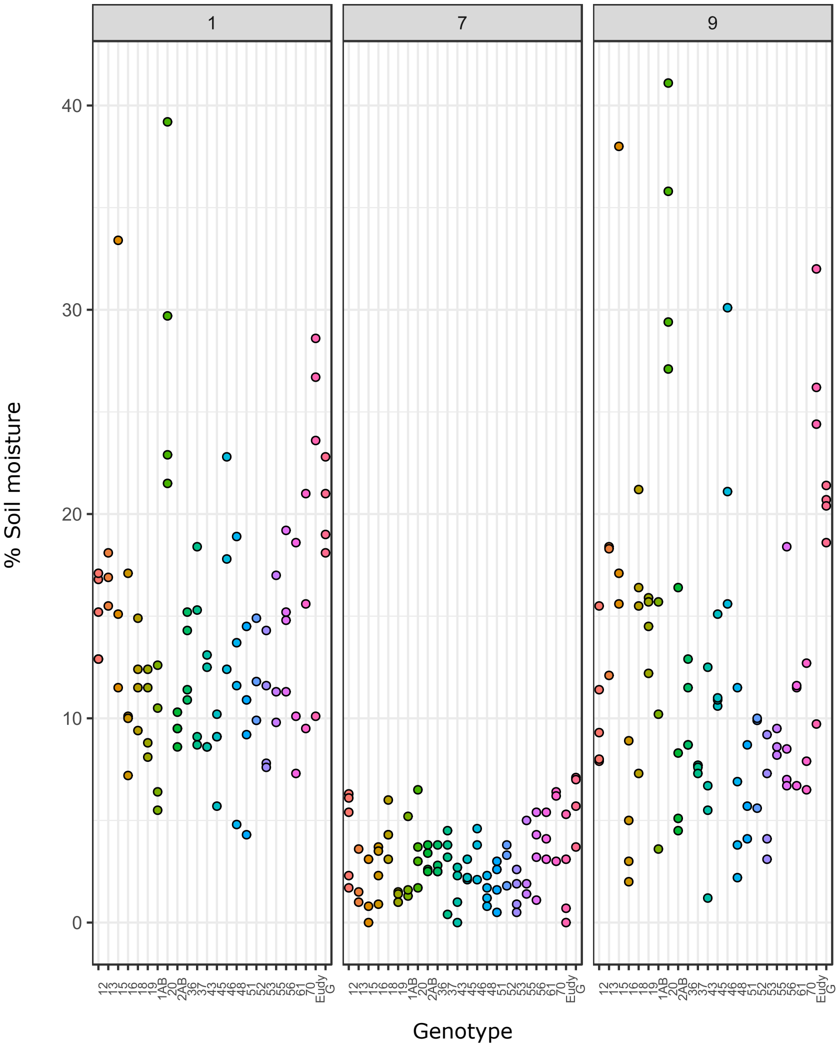


**Supplemental Figure 1** - Soil moisture measurements across replicate plants of each genotype. Showing only *Y. gloriosa* individuals measured as a part of this study.
