## Supplementary Figure 2 for "Leaf anatomy is not correlated to CAM function in a C_3_+CAM hybrid species, *Yucca gloriosa*"

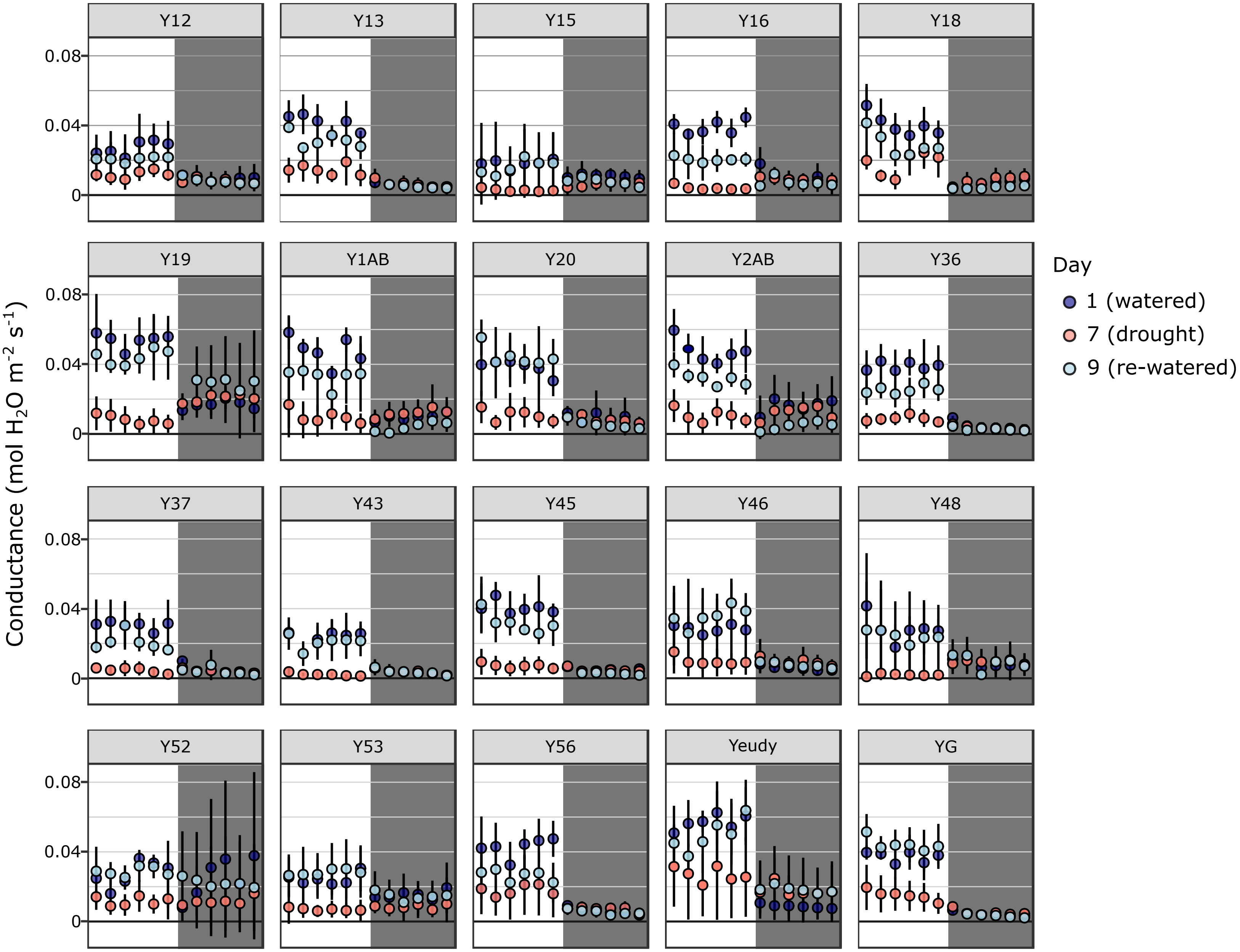


**Supplemental Figure 2** - Conductance measurements from Y. gloriosa genotypes measured every two hours over a 24 hour period, beginning 1 hour after the lights turned on (8 a.m.). White and grey backgrounds specify day and night timepoints, respectively. Mean and standard deviation are shown for day 1 (well watered), day 7 (drought), and day 9 (re-watered).
