## Supplementary Figure 3 for "Leaf anatomy is not correlated to CAM function in a C_3_+CAM hybrid species, *Yucca gloriosa*"

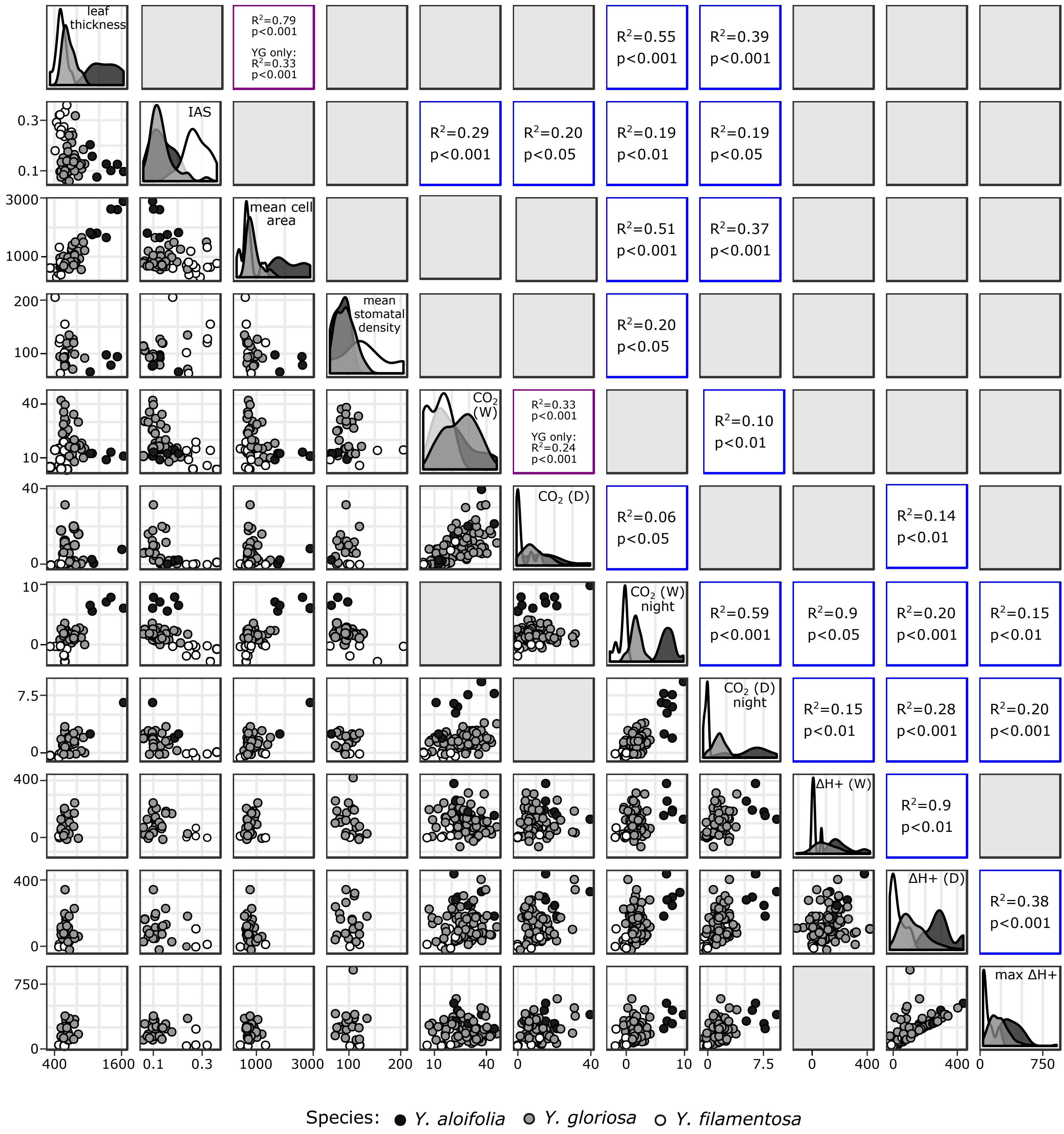


**Supplemental Figure 3** - Pairwise scatterplots for all traits on the lower diagonal, with density graphs on the diagonal and $R^2$ and p-value if significant on the upper diagonal. Purple outline indicates the two pairwise trait correlations that were significant in *Y. gloriosa*, and the corresponding $R^2$ and p-values are reported.
